# Bioinformatic analysis of the variability of Tax1 and Tax2 proteins of the human T-cell lymphotropic virus (HTLV)

**DOI:** 10.1101/2025.03.13.642961

**Authors:** Roberto Reinosa-Fernández, Francisco Javier Hernández-Walias

## Abstract

**Motivation:** The pathological differentiation in humans between HTLV-1 and HTLV-2 viruses is due, among other parameters, to the variability in a viral protein, Tax (Tax1 and Tax2 respectively), with the second virus being a candidate to interfere with HIV-1 infection, which has allowed us to assess the importance of further investigating Tax.

**Results:** Both the degree of conservation and the genomic and protein variability between Tax1 and Tax2 have been expressed in this work through bioinformatics tools. The knowledge about Tax2 related to the interfering capacity on HIV-1 compared to the absence of such a function in Tax1 is corroborated with our analyses, highlighting certain variations between both sequences that could explain the functional differences in pathogenicity that we show in this work.

## Introduction

HTLV is a retrovirus of the Deltaretrovirus genus, named by its English acronym (Human T cell Lymphotropic Virus). Its main target of action is the CD4+ T lymphocyte in the case of type 1 (HTLV-1), and the CD8+ T lymphocyte in the case of type 2 (HTLV-2) (Ijichi et al., 1992; Casoli et al., 2007; Biglione et al., 2013; Ciminale et al., 2014; Melamed et al., 2014). Additionally, it has been detected in monocytes, B lymphocytes, and dendritic cells (Biglione et al., 2013).

The complete genomes of HTLV-1 (Figure 1) and HTLV-2 (Figure 2) have certain similarities and differences in terms of gene representation in genomic and subgenomic fragments (Hulo et al., 2011). HTLV-1 consists of 8.5 kilobases (kb) and is made up of a genomic sequence and nine subgenomic sequences, of which the last two generate antisense transcripts. In the fifth subgenomic fragment, the **tax** gene is located, whose translated product is the Tax1 protein.

**Figure 1.**
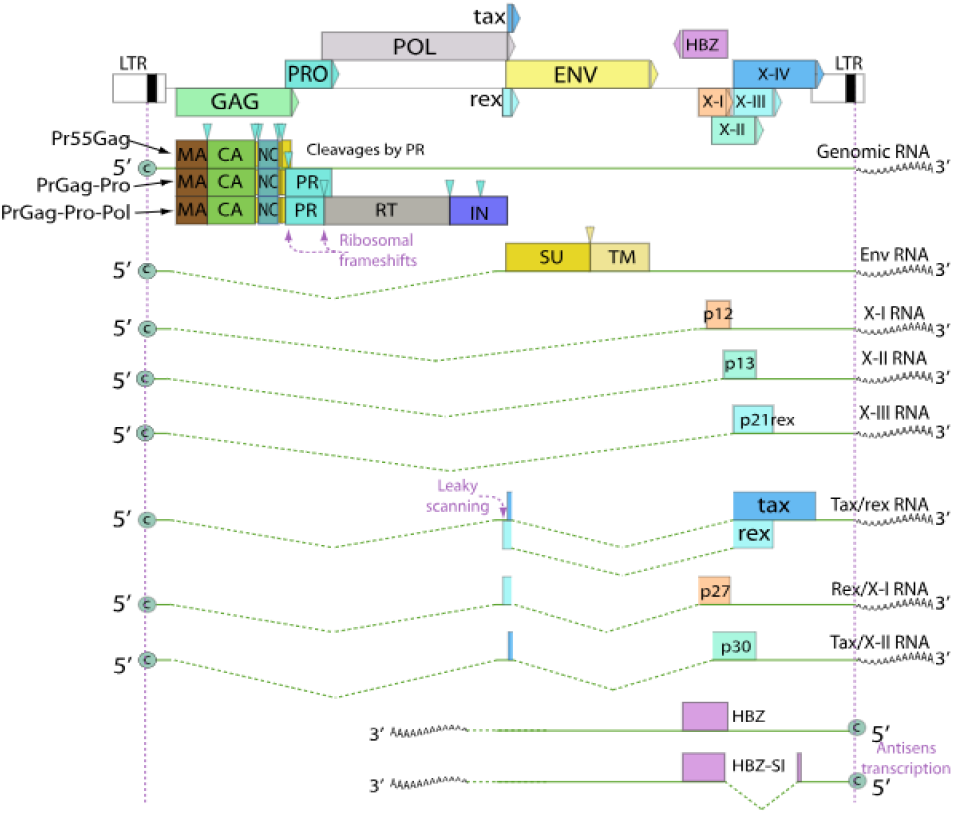
Genomic and subgenomic sequences of HTLV-1. (Retrieved from Hulo et al., 2011).

**Figure 2.**
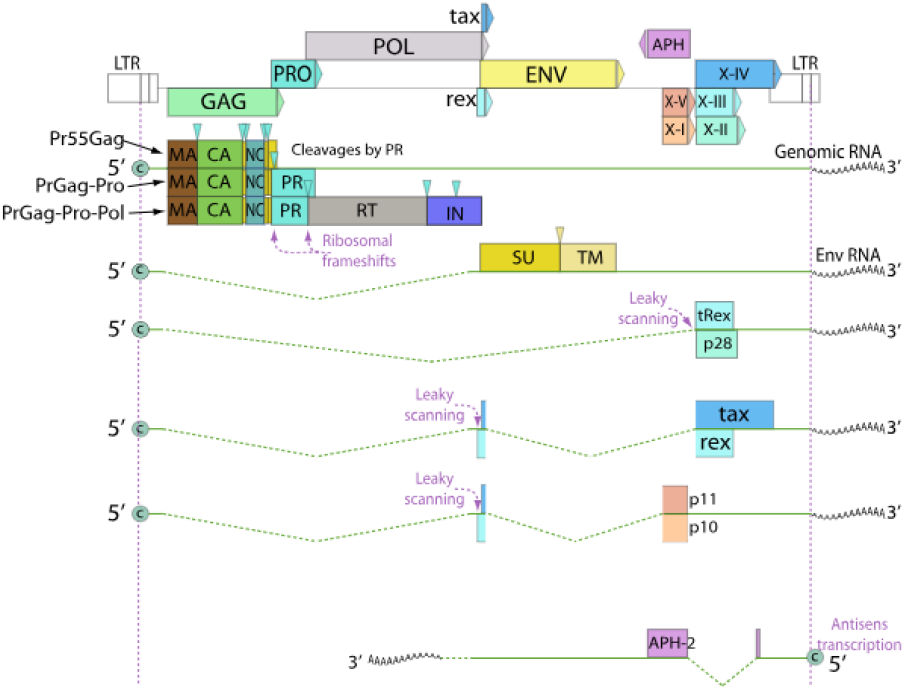
Genomic and subgenomic sequences of HTLV-2. (Retrieved from Hulo et al., 2011).

HTLV-2 consists of 8.9 kilobases (kb), however, it has a genomic fragment and five subgenomic ones, of which the last produces an antisense transcript. The **tax** gene is located in the third subgenomic fragment, whose translated product is the Tax2 protein.

The difference between the pathological implications of HTLV-1 and HTLV-2 in humans is very significant since the second does not seem to manifest specific diseases except for isolated cases of T lymphocyte neoplasms or neurodegenerative pathologies (Mahieux et al., 2000; Moreno et al., 2013) in the host, whereas the first is responsible for severe neurological diseases such as HAM (HTLV-1-associated myelopathy) and TSP (tropical spastic paraparesis) (Ciminale et al., 2014; Rivera-Caldón et al., 2017).

This work focuses on a series of bioinformatic analyses that compare Tax protein sequences in order to examine the variability of Tax1 and Tax2 and the differences between them.

## Materials and Methods

To carry out the bioinformatics analyses, the software MEGA and EpiMolBio have been used, as cited in articles such as Troyano-Hernáez et al., 2022. EpiMolBio was used to filter Tax1 and Tax2 sequences, perform alignments, calculate mutations of both proteins against the reference sequence, conservation studies, Wu-Kabat variability coefficient calculations, and consensus generation between Tax1 and Tax2. MEGA was used to manually eliminate artifact sequences.

EpiMolBio uses the color code shown below to display the percentages required according to the analysis. For this work, they indicate the frequency of appearance of the mentioned residues, whether the most conserved in conservation analyses, the mutated residue, or the consensus residue between both proteins.

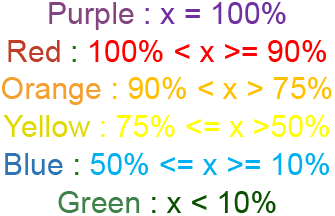

For both HTLV-1 and HTLV-2, all accessible protein sequences were downloaded from the NCBI Virus website (Virus NCBI, 2004). These were later filtered using the bioinformatics program EpiMolBio to obtain Tax1 and Tax2 from the full pool of sequences.

Once this was done, the sequences were aligned against the reference sequences provided by NCBI Virus, also using EpiMolBio software. Subsequently, artifact sequences were eliminated with the MEGA software (https://www.megasoftware.net/). The next step consisted of mutation analyses, consensus generation showing conservation, Wu-Kabat variability coefficient calculations, and consensus of consensuses generation using the EpiMolBio tool.

In total, 711 Tax1 sequences and 98 Tax2 protein sequences were used. It should be noted that a significant portion of these sequences are partial, however, EpiMolBio is capable of analyzing such sequences while ignoring gaps and unknown residues. The reference sequences used in this work are:

- NCBI Reference Sequence HTLV-1: NC_001436.1
- NCBI Reference Sequence HTLV-2: NC_001488.1

## Results

### 1. Descriptive analysis of mutations

A breakdown of the mutations analyzed in the studied samples of the Tax1 and Tax2 proteins has been made, in which the mutations with a frequency greater than or equal to 10% compared to the corresponding reference sequence are shown (Figure 3).

**Figure 3.**
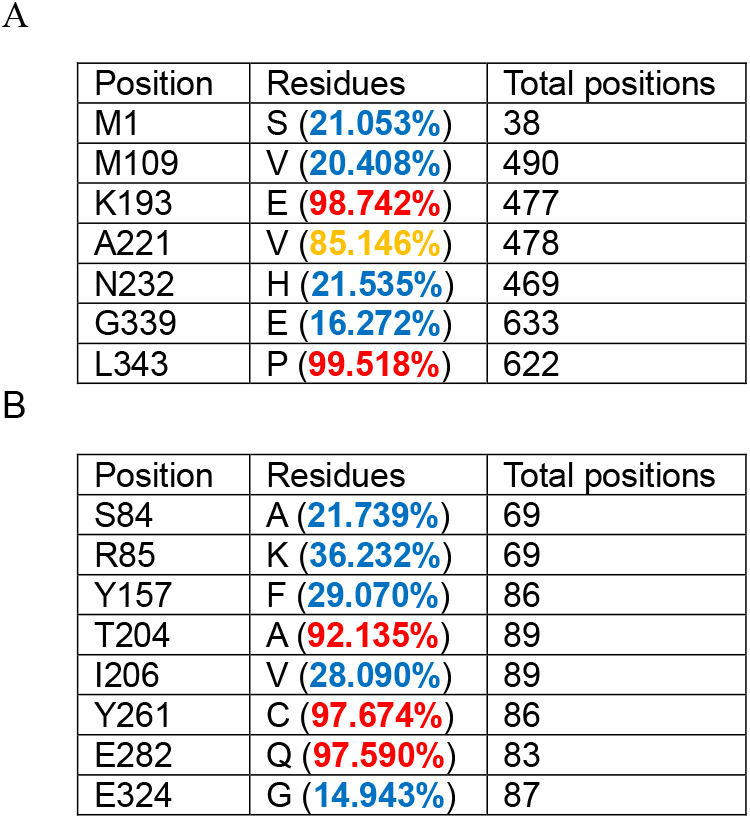
Table of equal or greater mutations frequency than 10% for Tax1 of HTLV-1 (A) and for Tax2 of HTLV-2 (B).

Here it can be seen that in Tax1, the mutations K193E, A221V, and L343P have a frequency of appearance greater than 85% in the introduced sequences compared to the reference sequence, just as for Tax2, we have that T204A, Y261C, and E282Q have a frequency of appearance greater than 90% compared to the reference sequence, which could be important for greater biological efficacy.

### 2. Variability in the residues of Tax1 and Tax2

An analysis of the conservation of Tax1 and Tax2 is shown through a consensus marked with the color code of the EpiMolBio program corresponding to the differences in the frequencies of appearance of the residues:

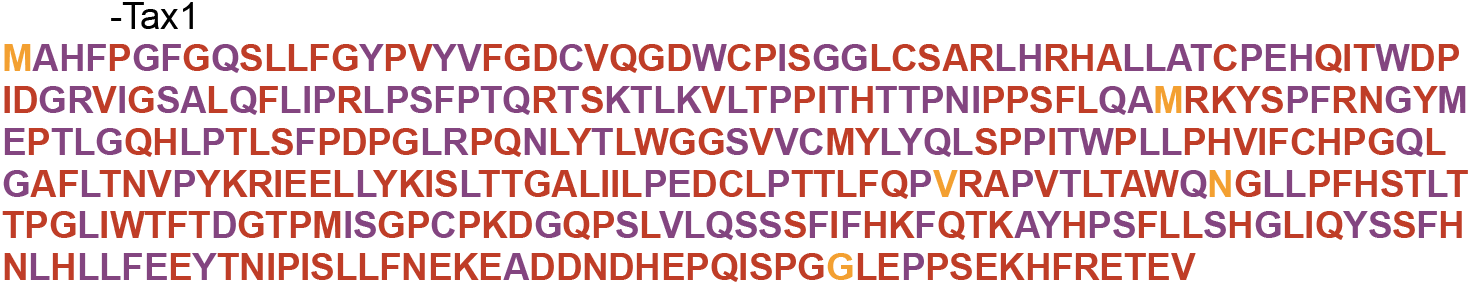

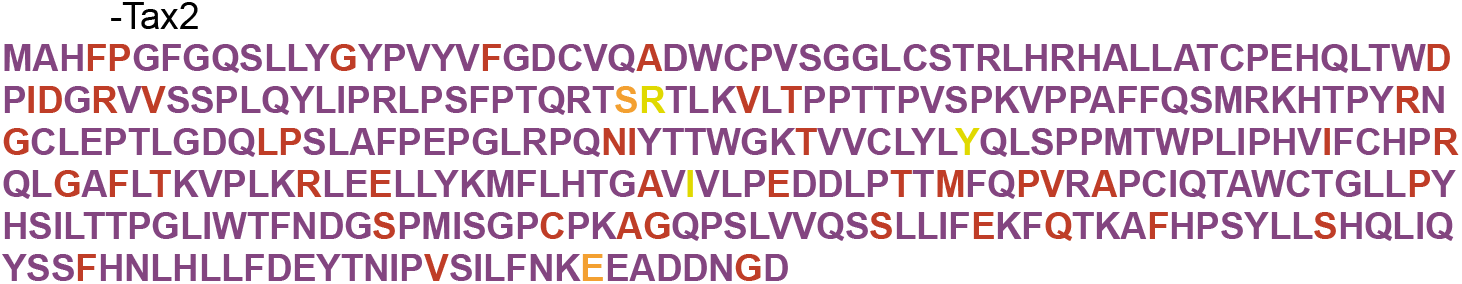

These results show that both proteins are highly conserved since most of the residues of the consensuses appear colored in red or purple. The purple residue means that 100% of the sequences had that residue at that position, and red means a percentage equal to or greater than 90%. Some exceptions can be observed, but generally, a very high conservation is observed.

To evaluate the changes in the residues of Tax1 and Tax2 in all the analyzed sequences, the Wu-Kabat variability coefficient has been measured (Figure 4 and Figure 5).

**Figure 4.**
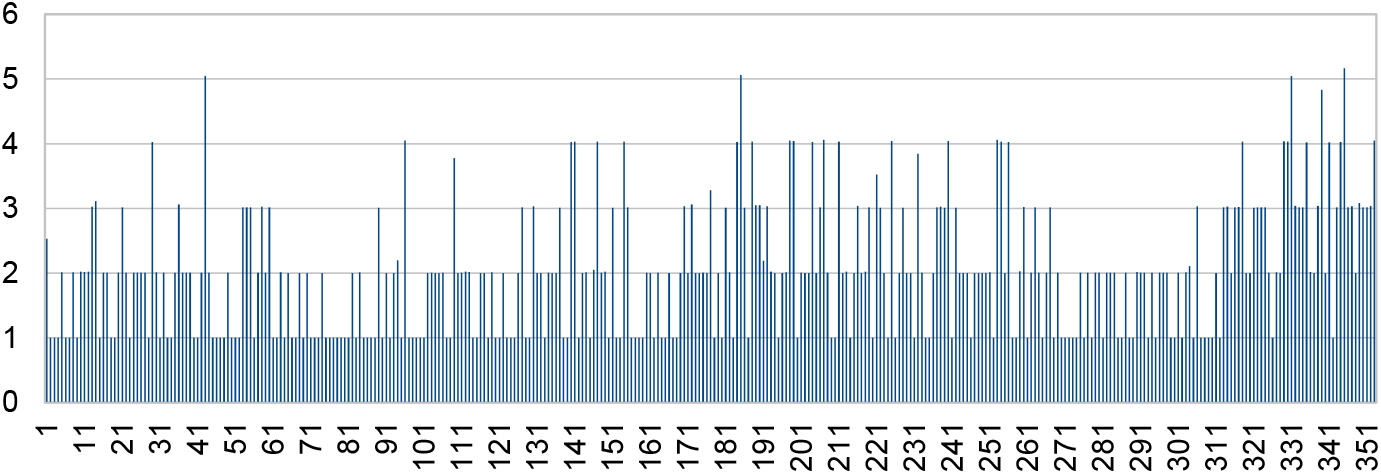
Graph of the Wu-Kabat variability coefficient for each residue of the Tax1 protein.

**Figure 5.**
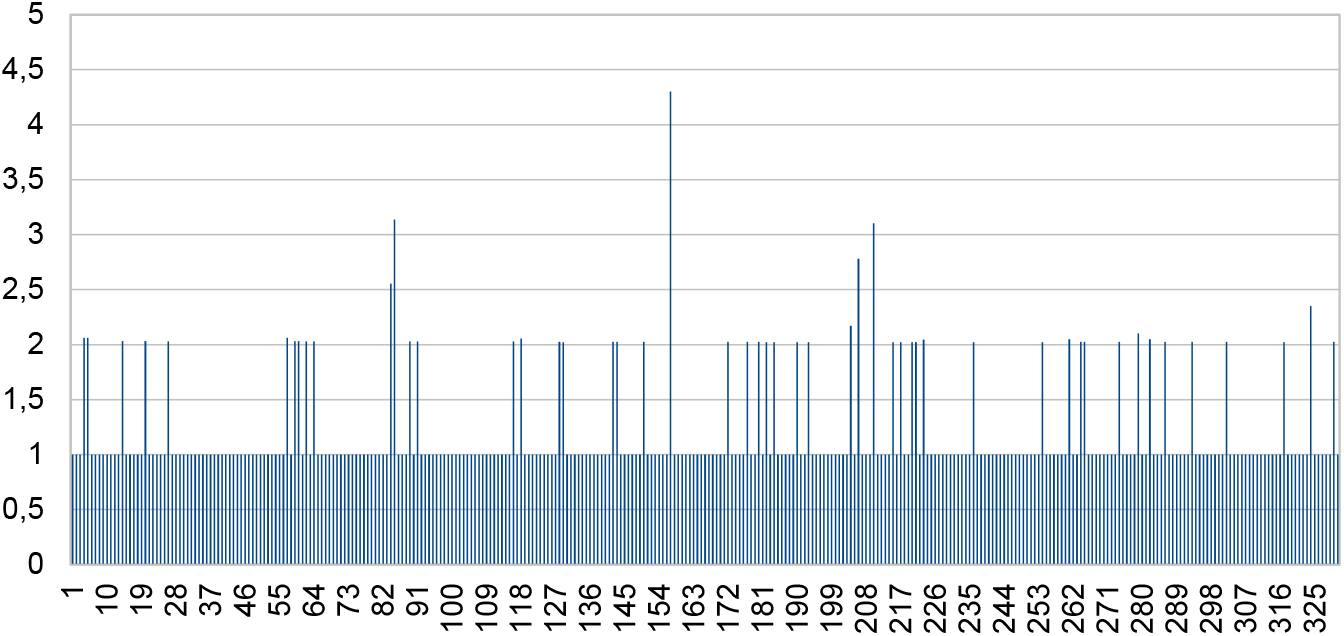
Graph of the Wu-Kabat variability coefficient for each residue of the Tax2 protein.

### 3. Generation of consensus sequence between Tax1 and Tax2

A consensus of consensuses of Tax1 and Tax2 has been generated to see common and variable regions. By aligning the protein sequences of Tax1 and Tax2, residue 331 is the last common one, that is, from residue 332 onwards, there are no residues in the Tax2 sequence, so gaps have been introduced in the alignment with Tax1, which goes up to residue 353 (Figure 6). Therefore, the residues that appear from position 331 onwards have been excluded from the analysis, as it would only be a consensus of Tax1.

-Consensus of consensuses TAX-1 and TAX-2:

**Figure 6.**
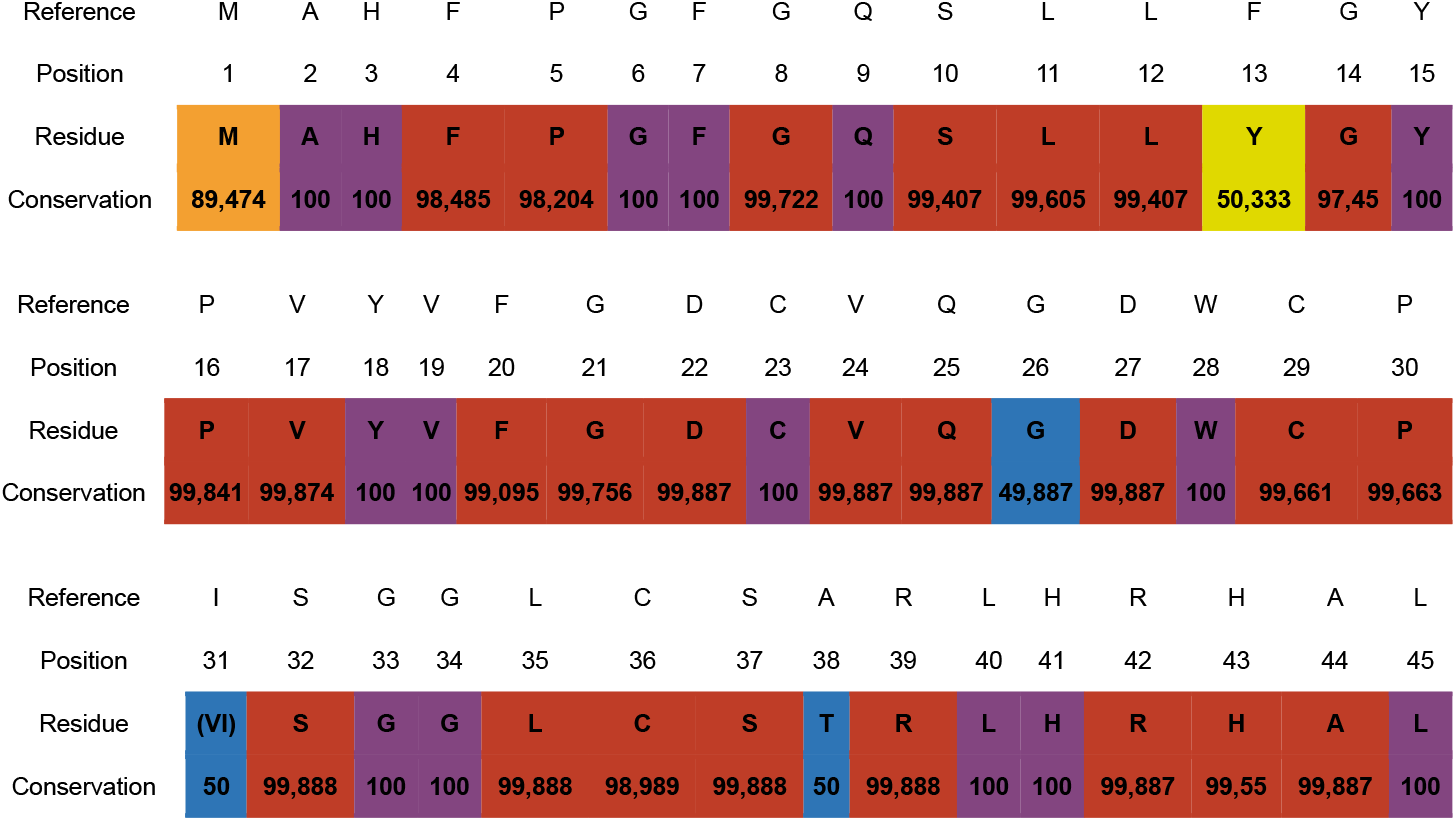
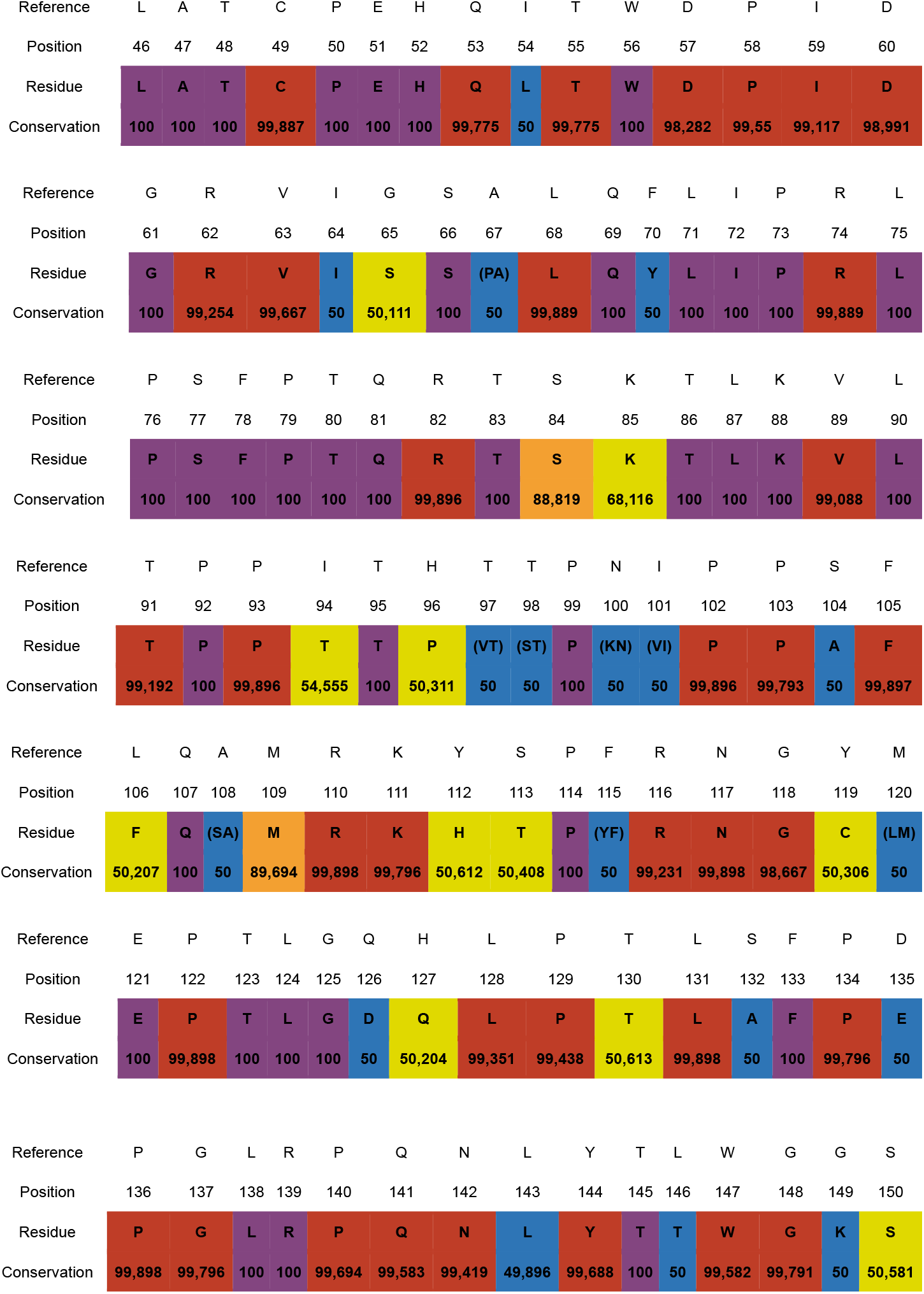
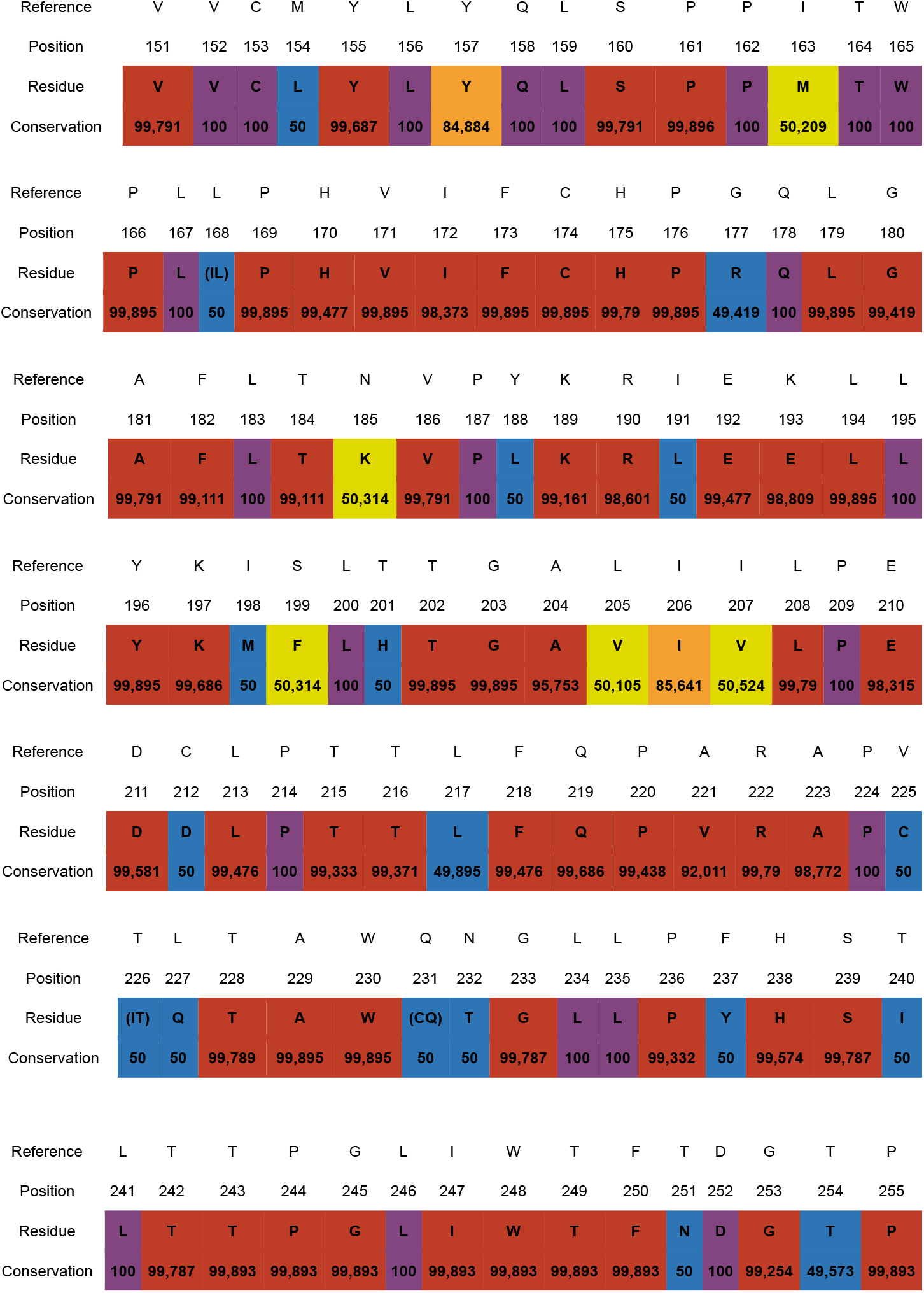
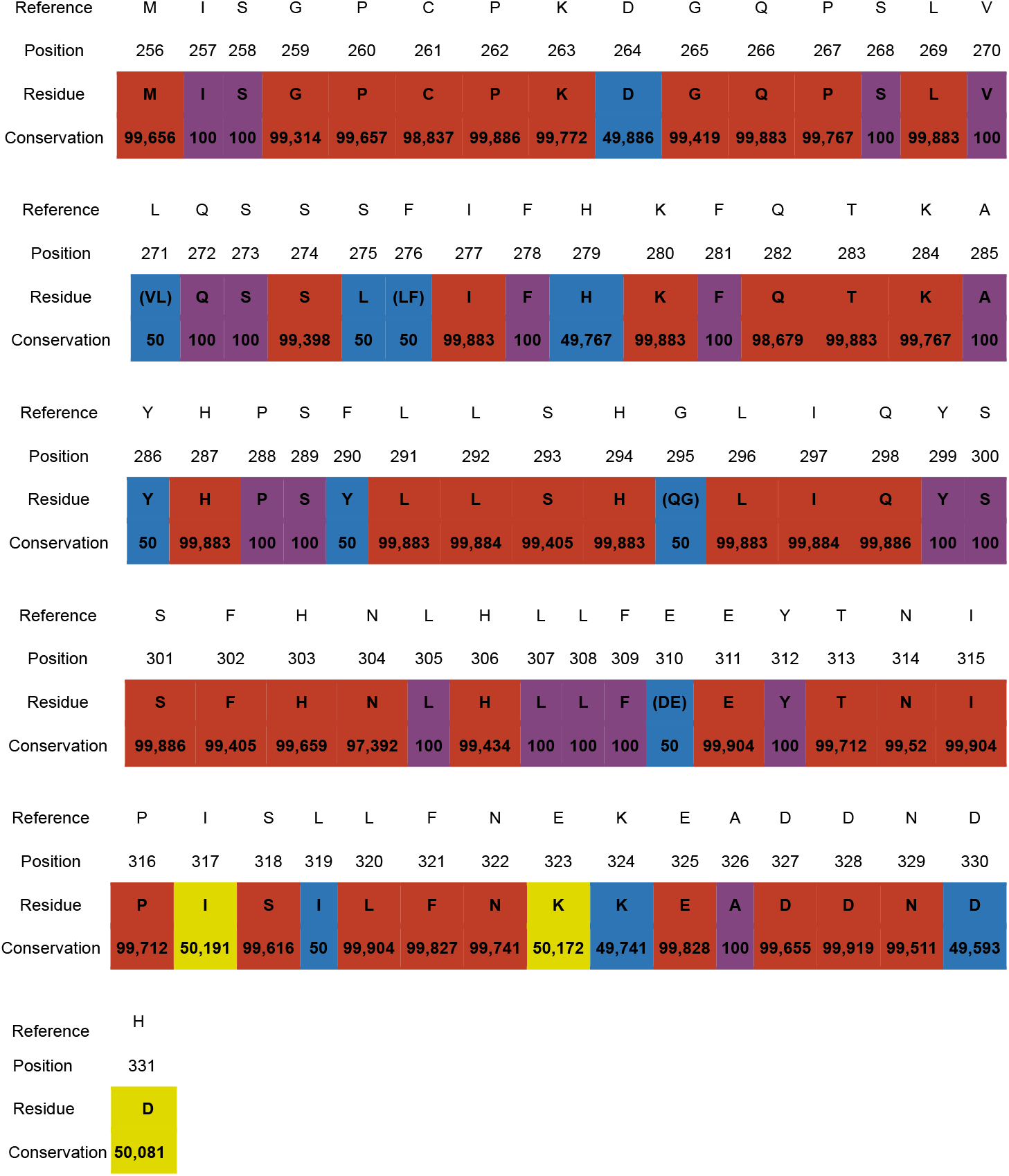
Consensus table of consensuses between Tax1 and Tax2.

In this table, the consensus of consensuses between Tax1 and Tax2 is shown, where it can be seen that there are regions with high similarity between both proteins, showing conserved regions between both. However, variable regions are also shown.

## Discussion

Considering regions 1, 13, 16, 43, 45, 49, and 54 studied in Ross et al., 1997 corresponding to Tax1, and taking into account the consensus between Tax1 and Tax2, there are differences between these two proteins since the conservation frequency of the consensus of consensuses is not too high in positions 1, 13, and 54 of our study, which implies a difference between both proteins. This has been observed in our results in region 1 where M1S is mutated in 21.053% in Tax1, indicating that there is a mutation in this position as important as mentioned in Ross et al., 1997. Between Tax1 and Tax2, we have detected differences in positions 13 and 54, which, despite being highly conserved in Tax1 and Tax2, are different between both proteins. All include activation domains at the amino- and carboxyl-terminal ends.

There are dissimilar zones between Tax1 and Tax2, marked in yellow and blue, which would correspond to a frequency less than or equal to 75% in our work (Figure 6) that could provide an explanation for the protective difference between HTLV-1 and HTLV-2 when considering the expression of HIV-1 blocking cytokines. According to the studies of Shoji et al., 2009, there is a pathogenic difference between Tax1 and Tax2. Although it does not refer to the action on HIV-1 infection, it does take into account the different pathogenic implications of both proteins in a lymphoid line. This research group shows that between Tax1 and Tax2, the former has the ability to transform a mouse lymphoid line (CTLL-2) with IL-2 dependence into independent growth, making this transformation much more potent compared to the latter.

In our study, we have seen that there is a high conservation in the Tax1 and Tax2 proteins, but with certain differences between them. This differentiation contributes to some extent to the different genetic pattern between HTLV-1 and HTLV-2, corroborated in other studies where it is shown that there is a 70% similarity of their nucleotides (Mahieux et al., 2000; Martínez et al., 2019). This incomplete genetic similarity may provide an explanation regarding differences in clinical manifestations (Ciminale et al., 2014; Martínez et al., 2019).

The difference between the sequences of Tax1 and Tax2 that we have observed could indicate that, since these Tax proteins induce the production of MIP-1α, in an HTLV-2 infection higher levels of this cytokine are generated by CD8+ T lymphocytes, and this contributes to its binding to the CCR5 coreceptor of CD4+ T lymphocytes, blocking their effective interaction with HIV-1 (Gordon et al., 2010). However, much more exhaustive studies are needed to affirm this since Tax1 also allows the release of MIP-1α, and its difference with Tax2 could potentially contribute to the differential release of this cytokine by both viruses, and it is probably not only due to the cell type.

Specifically, in the case of the Tax2 protein, there is evidence of its relationship with an inhibition role of apoptosis (Melamed et al., 2014), which could be corroborated by the high levels of CD8+ T lymphocytes that exist in individuals co-infected with HIV-1 and HTLV-2 compared to samples from HIV-1+ individuals without HTLV-2 infection (Hernández-Walias, 2022).

The co-infection relationship between HIV-1+ individuals and HTLV-2 is known, which, as previously mentioned, implies that the latter can block the infection of the former in CD4+ T lymphocytes through the action of the Tax2 protein from HTLV-2 that induces the expression of cytokines such as IFN or MIP-1α from CD8+ T lymphocytes (Pilotti et al., 2007).

As in HTLV-2 infection, in the case of HTLV-1, the release of MIP-1α also occurs, in addition to other cytokines such as IP-10. These produced levels have been significantly higher in HTLV-1 infection than in patients not infected by this virus (Yamamoto et al., 2004).

## Conclusions

The complete analysis of the protein sequences of Tax1 and Tax2 can provide relevant information to evaluate whether the difference in the pathogenicity of HIV-1 by both viruses is influenced by these proteins. In this study, we aim to compare both proteins to see if there is a difference in the production of MIP-1α, knowing that both viruses produce them.

The comparison of Tax1 and Tax2 indicates potential changes in the three-dimensional structure of both proteins, which may explain their differential effect on the production of MIP-1α, and this could suggest that potentially not only the cell type is responsible for the different production of MIP-1α but also the aforementioned different three-dimensional structure.

